# *SlNAP2* promoted fruit ripening by directly binding the *ACS2* promoter and interacting with *EIL3*

**DOI:** 10.1101/2024.07.15.603576

**Authors:** Xuetong Wu, Hua Fang, Dengjing Huang, Xuemei Hou, Yali Qiao, Changxia Li, Ailing Li, Yi Huang, Zhongxing Zhang, Zhiya Liu, Yayu Liu, Weibiao Liao

**Author notes:** **Corresponding author**: Weibiao Liao. The author responsible for distribution of materials integral to the findings presented in this article in accordance with the policy described in the Instructions for Authors (https://academic.oup.com/plphys/pages/General-Instructions) is Weibiao Liao. **Author contributions**. X.W. and W.L. designed the experiments, analyzed the data and wrote the manuscript. X.W., H.F., D.H., X.H., Y.Q., A.L., and Y.H. performed most of the experiments. assisted by Z.L., Z.Z., and Y.L. All authors, especially X.W. and W.L. commented on the manuscript. All authors read and approved the manuscript.

## Abstract

The ripening process of tomato fruit is affected by a variety of environmental factors and genetic regulators. NAC transcription factors (TFs) function in a multitude of biological processes, while the current knowledge on the participation of NAC TFs in the regulatory network of fruit ripening is relatively limited. In this study, we isolated a NAC TFs, NAP2, which acts as a positive transcription activator in tomato fruit ripening. We also observed a notable delay in the ripening process of *SlNAP2* silenced and knockout mutant fruit. In particular, ethylene production was obviously inhibited in *SlNAP2* mutant fruit. Y1H and DLR assays showed that SlNAP2 directly binds to the promoter of *SlACS2* and activates its transcriptional activity. Furthermore, SlNAP2 and SlEIL3 physically interaction was demonstrated by yeast two-hybrid (Y2H), luciferase complementation (LUC), bimolecular fluorescence complementation (BiFC) and coimmunoprecipitation analysis (CoIP) assays. Meanwhile, the pigment content, firmness and the transcript levels of genes associated with carotenoid and chlorophyll metabolism and cell wall metabolism were also potentially affected by the *SlNAP2* deletion; however, it has remained unclear whether these genes are also directly regulated by *SlNAP2*. Therefore, our findings indicate that SlNAP2 directly binds to *SlACS2* promoter to activate its expression and promote ethylene generation, which in turn interacts with EIL3 to enhance the function of ethylene in tomato fruit ripening. Collectively, our data contribute to understanding the interaction of NAC TFs and ethylene in tomato fruit ripening, thereby enhancing our knowledge of the ripening regulatory network that governs tomato fruit maturation.

## Introduction

Tomato is favoured by consumers for its unique flavour and high nutritional value, and has high economic benefits in the present development of facility horticulture (Tieman et al., 2017). Tomato is rich in nutrients, including vitamin C, soluble sugar, soluble protein, lycopene and so on. Tomato has become the best industry crop for market demand (Yan et al., 2021). Additionally, tomato fruit, as a typical model, are widely used to explore the ripening of climacteric fruit (Feder et al., 2020). The process of tomato fruit ripening is a genetically maneuvered and intricate process that results in significant changes in the physical characteristics of the fruit, including flesh color, flavor, texture, and nutritional ingredient (Martel et al., 2011). Fruit ripening is influenced by hormonal, environmental and genetic regulators which play a decisive role in the maturation signal network (Karlova et al., 2014; Ji et al., 2021). The morphological changes observed during fruit ripening are the consequence of the simultaneous activation of various biochemical and genetic pathways (Shan et al., 2012; Gao et al., 2022). To enhance and enrich the regulatory mechanism and network model of climacteric fruit ripening, it is crucial to explore and identify more transcription factors (TFs), their regulated target genes, and interacting proteins.

Researchers propose that NAC TFs form a distinct family in plants (Feng et al., 2023). NAC proteins are generally no more than 400 amino acids in length. The N-terminus contains a highly conserved NAC structural domain, and the C-terminus represents a flexible transcriptional regulatory region with both transcriptional activating and transcriptional repressing activities, but the region also exhibits intense complexity (Zhu et al., 2014). The NAC TFs family in plants is substantial and multifunctional, and 101 NAC genes in tomato have been identified in the Plant Transcription Factor Database (http://planttfdb.cbi.pku.edu.cn/), which are participated in various processes and signaling networks related to plant growth, development, senescence, ripening and stress tolerance (Zhang et al., 2011). Based on our previous research that SlNAP1, characterized as a member of the NAC TFs, accelerated fruit ripening, and target genes (*SlGA2ox1* and *SlGA2ox5*) directly activated by SlNAP1 (SlGA2ox1 and SlGA2ox5) were identified (Li et al., 2024). Similarly, model fruit tomato ripening is also dependent on the NAC TFs, *NOR-like 1*, which directly targeted the promoters of cell wall metabolism associated genes (*SlPG2a、SlPL、SlCEL2* and *SlEXP1*), chlorophyll degradation related genes (*SlGgpps2、SlSGR1*), and ethylene synthesis genes (*SlACS2、SlACS4*) to stimulate their expression (Gao et al., 2022). *NOR-like 1 s*erves as a crucial player in the mature network, acting as a positive activator in tomato. Meanwhile, researchers also demonstrated that *SlNAC9* knockout caused delayed fruit ripening and directly regulated *SGR1*, *DXS2*, *PSY1* and *CrtR-b2* to suppress the relative expression of carotenoid metabolism-associated genes (Feng et al., 2023). However, *SlNAC1* was identified to suppress fruit ripening by regulating the promoter regions of ethylene and lycopene synthesis-associated genes (Ma et al., 2014). Thus, the engagement of NAC TFs in the modulation of fruit ripening varies and the elaborate functioning is not well developed.

Endogenous ethylene generation is a marker substance in climacteric fruit ripening (Bleecker, 2000). The ingredient participated in ethylene biosynthesis and signaling network, including ethylene-responsive factors (SlERFs), 1-aminocyclopropane-1-carboxylic acid oxidases (SlACOs), 1-aminocyclopropane-1-carboxylic acid synthases (SlACSs), and ethylene-insensitive factors (EIL3s), modulate fruit ripening rate and quality through diverse mechanisms (Huang et al., 2022; Liu et al., 2015). Although each component may overlap in function, each also assumes a unique role, and together they coordinate and control the ripening process of tomato fruits (Baranov and Timerbaev, 2024). An ethylene receptors (ETRs) family gene, *SlETR7*, was newly identified, and overexpression of *SlETR7* caused sooner flowering, poorer plants, and smaller fruits, and the *sletr7* knockout mutant was able to produce more ethylene at Br stage (Chen et al., 2020). The study reported that a combined deletion of *SlACS2* and *SlACS4* blocked the generation of the ethylene burst which are necessary required for the normal process of fruit ripening in tomato (Hoogstrate et al., 2014). Gao et al. (2021) characterized *SlNAM1*, a NAC TFs, which directly bound to ethylene biosynthesis genes (*SlACS4, SlACS2*) and positively influenced fruit ripening of tomato. Interesting, researchers suggested that NAC TFs is also responsive to ethylene induction (He et al., 2005; Xia et al., 2010) and act as a TFs downstream of the EIN2 in conjunction with EIN3 (Kim et al., 2009; Al-Daoud, 2011). MaNAC1/MaNAC2 in banana engaged in physical interaction with EIN3-like protein, designated MaEIL5, which together mediated banana ripening (Greco et al., 2012). Therefore, NAC TFs and ethylene signalling synergistically coordinate the fruit ripening process and ensure that this process occurs in a timely and efficient manner. This coordination is essential for proper fruit development.

Although the joint modulation of tomato fruit ripening by ethylene and NAC TFs has been reported in the literatures, it is only relatively superficially and shallowly understood. Due to the magnitude and richness of the tomato genome, a well-developed and comprehensive regulatory signaling network has yet to be developed. Therefore, it is beneficial to explore more interactions mechanisms of NAC TFs related to ethylene synthesis and signaling components in order to better improve the fruit ripening signaling network. In this study, we knocked out the *SlNAP2* gene using CRISPR/Cas9 technology and focused on how *SlNAP2* influences ethylene synthesis and signaling components in tomato, as well as the target genes of *SlNAP2* that regulate ethylene synthesis. This study provides some new information for the establishment of a regulatory network for tomato ripening.

## Results

### Isolation and expression patterns of *SlNAP2*

The *SlNAP2* gene (NM_001365397.1) (1226bp) is situated in chromosome 4 and contains three exons and two introns. Amio acid multiple sequence alignments demonstrated that SlNAP2 is a member of NAC TF family and contains the highly conserved NAC domain (Fig. 1A). The full-length CDS of *SlNAP2* spans 828 bp, encoding the SlNAP2 protein consisting of 279 amino acids. Phylogenetic showed that *SlNAP2* and *SlNAP1* are closely related NAC TFs in tomato, and homologous to *AtNAP* (Supplemental Fig. S1). The CDS of *NAP2* was amplified and isolated from ’Mirco Tom’. Subsequently, the expression information *SlNAP2* gene was analyzed in tomato. We harvested the roots, stems, leaves, flowers and fruits (MG, Br and Br + 10) in tomato and then investigated the transcriptional expression pattern of *SlNAP2* by qRT-PCR assay. The *SlNAP2* transcripts were detected in different tissues and has the highest expression abundance in fruits at Br stage (Fig. 1B). Additionally, to confirm the spatial expression properties of the SlNAP2 protein, the CDS of *SlNAP2* was fused in frame with the *35S:: GFP* (Supplemental Fig. S1). The GFP fluorescence from the *35S::GFP-SlNAP2* construct is localized to the cell membrane, whereas the fluorescence from the only GFP-carrying construct is distributed across the cell membrane and nucleus (Supplemental Fig. S1). The transcriptional activity investigation of SlNAP2 was validated by Y1H assay. Yeast cells containing the full-length structural domain of SlNAP2 exhibited robust growth in SD/-Trp-His-Ade medium, and the results suggest that SlNAP2 possesses individual transcriptional activity (Fig. 1C and Supplemental Fig. S1).

**Figure 1.**
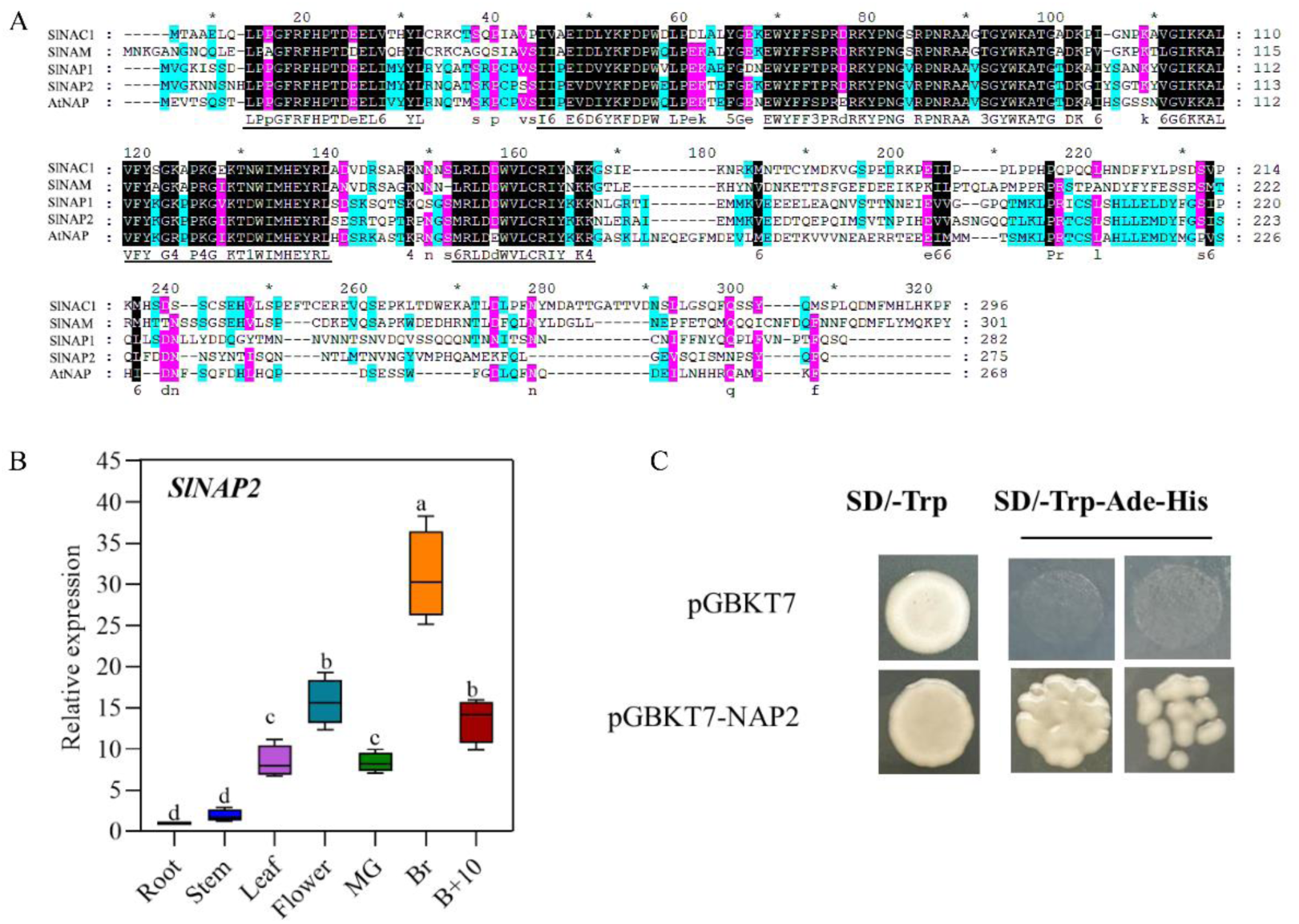
Characterization and isolation of the *SlNAP2* gene. **A)** Amino acid sequence alignment of SlNAC1, SlNAM, AtNAP, SlNAP1, and SlNAP2 proteins. Identical amino acids in all five proteins are highlighted with a black background. The five conserved subdomains in the NAC domain are marked in black above the sequences. **B)** Expression pattern of *SlNAP2* in different organs of tomato. **C)** Transcriptional activation assay of *SlNAP2* in Y2H yeast cells. *SlNAP2* exhibited transcriptional activation ability. The different lowercase letters signify significant differences with a *p* <0.05, as evaluated by Duncan’s test.

### *SlNAP2* accelerates the tomato fruit ripening

To preliminary assess the role of *SlNAP2* in ripening of tomato fruit, we obtained stable functional deletion *slnap2* mutant (*slnap2-3* and *slnap2-6*) using the CRISPR/Cas9 gene editing system. A specific single guide RNA (sgRNA) was devised to specifically target in the first exon of *SlNAP2*. The amplified sgRNA sequence was inserted into a vector containing the sgRNA and Cas9 expression cassette. Subsequently, the construct was introduced into WT tomato cv. ‘MicroTom’ by infecting leaf explants with *Agrobacterium*. Transgenic lines obtained were genotyped by direct sequencing of PCR reaction solution of genomic DNA surrounding the target site. The two mutants contained the same homozygous 379 th base deletion, and the same homozygous addition of one base between bases 378 th and 379 th in the first exon of *SlNAP2*, respectively (Fig. 2A). Both the base 379 deletion and the base 379 insertion mutations are anticipated to result in a premature stop codon in the first exon of *SlNAP2*. We referred to the mutant with the deletion at base 379 th as *slnap2-3* and with the addition at base 379 th as *slnap2-6* (Supplemental Fig. S2). In addition, we gained *SlNAP2*-silenced line (TRV1+TRV2-*NAP2*). Subsequently, gene expression of *SlNAP2* in tomato fruits at MG and Br was analyzed by qRT-PCR. Significant inhibition of *SlNAP2* gene expression was observed in both stages, reaching a silencing efficiency of more than 70% (Fig. 2B). Fruit ripening in the *SlNAP2*-silenced lines, *slnap2-3* and *slnap2-6* mutants was significantly delayed, compared with fruit ripening in WT (Fig. 2C). We compared fruit of WT, the Sl*NAP2* silencing lines and the *slnap2* mutants at four stages: mature green (MG), breaker stage (Br), 5 days after breaker stage (Br+5) and red and ripening (Re). The WT fruits turned red, while the fruit from the silencing lines and the mutants remained at the Br stage. Although the suppressive effectiveness of the silencing lines (TRV1+TRV2-*NAP2*) was not as pronounced as the *slnap2-3* and *slnap2-6* mutants, their results were consistently similar, demonstrating that *SlNAP2* is critical for tomato ripening fruits. To precisely investigate the variation in *slnap2* mutant fruit color, we also measured L*, a*, and b* in MG and Br fruit. The results show that in the *slnap2-3* and *slnap2-6* mutants a* and b* were prominently lower than in the WT plants, and a* and b* in the silenced lines although higher than in the mutants, remained lower than in WT plants (Fig. 2E). Conversely, L* exhibited the exact opposite trend. We also observed that L*, a*, and b* at MG stage did not differ significantly among the three lines (Fig. 2D). Consequently, *SlNAP2* plays a positive activator role in ripening of tomato fruit, and the deletion of *SlNAP2* significantly delays tomato fruit ripening.

**Figure 2.**
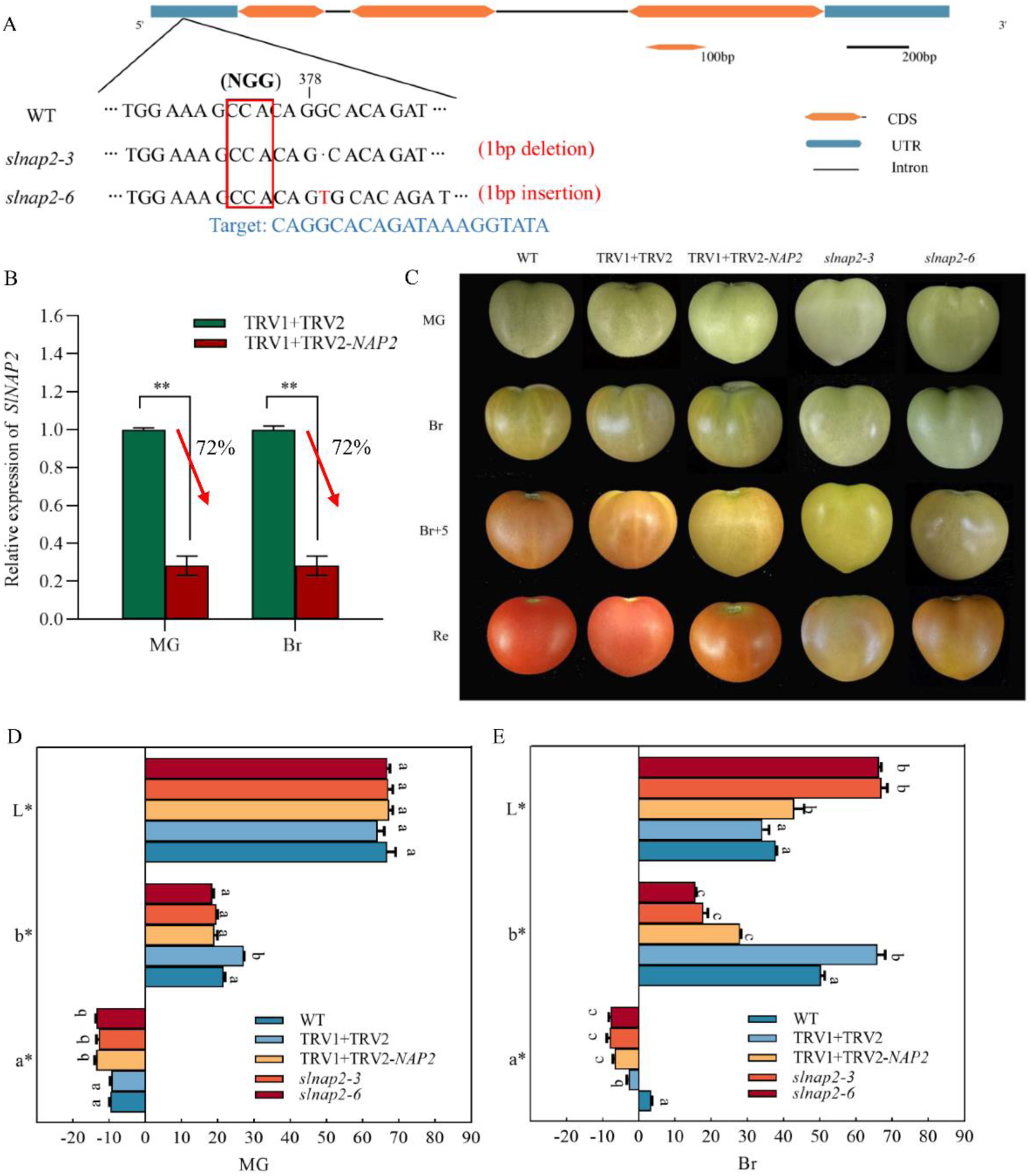
The phenotype of tomato fruit ripening in the three lines. **A)** Gene editing identification of *slnap2-3* and slnap2-6 mutants. Red letters indicate edit site and edit type. Blue letters indicate the target sequence. **B)** qRT-PCR of VIGS-induced *SlNAP2* gene silencing at MG and Br stages. The gene silencing efficiency reached 72%. **C)** Phenotype of tomato fruit in the three lines. **D)** L*, a* and b* in the three lines at MG and Br stages. The different lowercase letters signify significant differences with a *p* <0.05, as evaluated by Duncan’s test.

### *SlNAP2* affects chlorophyll degradation and carotenoid synthesis in tomato fruit

To gain understanding into the inhibition of fruit ripening by the deletion of *SlNAP2*, we measured pigment content in fruit of three lines. The *slnap2-3* and *slnap2-6* mutant fruit had marked elevation in chlorophyll a and chlorophyll b contents at MG stage compared to WT fruit (Fig. 3A). Similarly, chlorophyll a content was considerably greater in *slnap2-3* and *slnap2-6* mutant fruits at Br than in WT and TRV1+TRV2 plants (Fig. 3B). Carotenoid content was dramatically decreased in the mutants and silencing lines than in the WT plants, whether at MG and Br stage. However, chlorophyll b content in *slnap2* mutant fruits at Br was not appreciably different in WT and silenced lines. To further understand whether *slnap2* mutants delay ripening by regulating pigment synthesis in fruit, we randomly selected five chlorophyll degradation-associated genes and carotenoid synthesis-associated genes, respectively, and investigated their expression levels in WT plants, mutants, and silencing lines. The results revealed that the expression abundance of genes associated with chlorophyll degradation and carotenoid synthesis in both MG and Br stage mutant fruits were substantially remarkably lower than in WT fruit. Compared with WT plants, *PAO* expression level in *slnap2-3* and *slnap2-6* mutant fruits was down-regulated by 90.9% and 78.9% at MG stage, respectively (Fig. 3C). The transcript levels of *NYC*, *SGR*, *PAO*, *PPH*, and *RCCR* were all downregulated by more than 76.8% in mutant fruits in comparison to WT plants at Br stage (Fig. 3D). At MG stage, the transcript levels of carotenoid synthesis genes were similarly reduced in mutant and silencing line fruits (Fig. 3E). Notably, the *PSY1*, *GGPP*, *CYHB2*, *LYCE*, and *ZEP* expression levels were also substantially downregulated in *SlNAP2* mutant fruits at the Br stage (Fig. 3F). This implied that the stage at which the *slnap2* mutant had the greatest beneficial influence on fruit ripening was at breaker stage. The *SlNAP2* knockout might affect the chlorophyll degradation and carotenoid synthesis to further delaying the fruit ripening.

**Figure 3.**
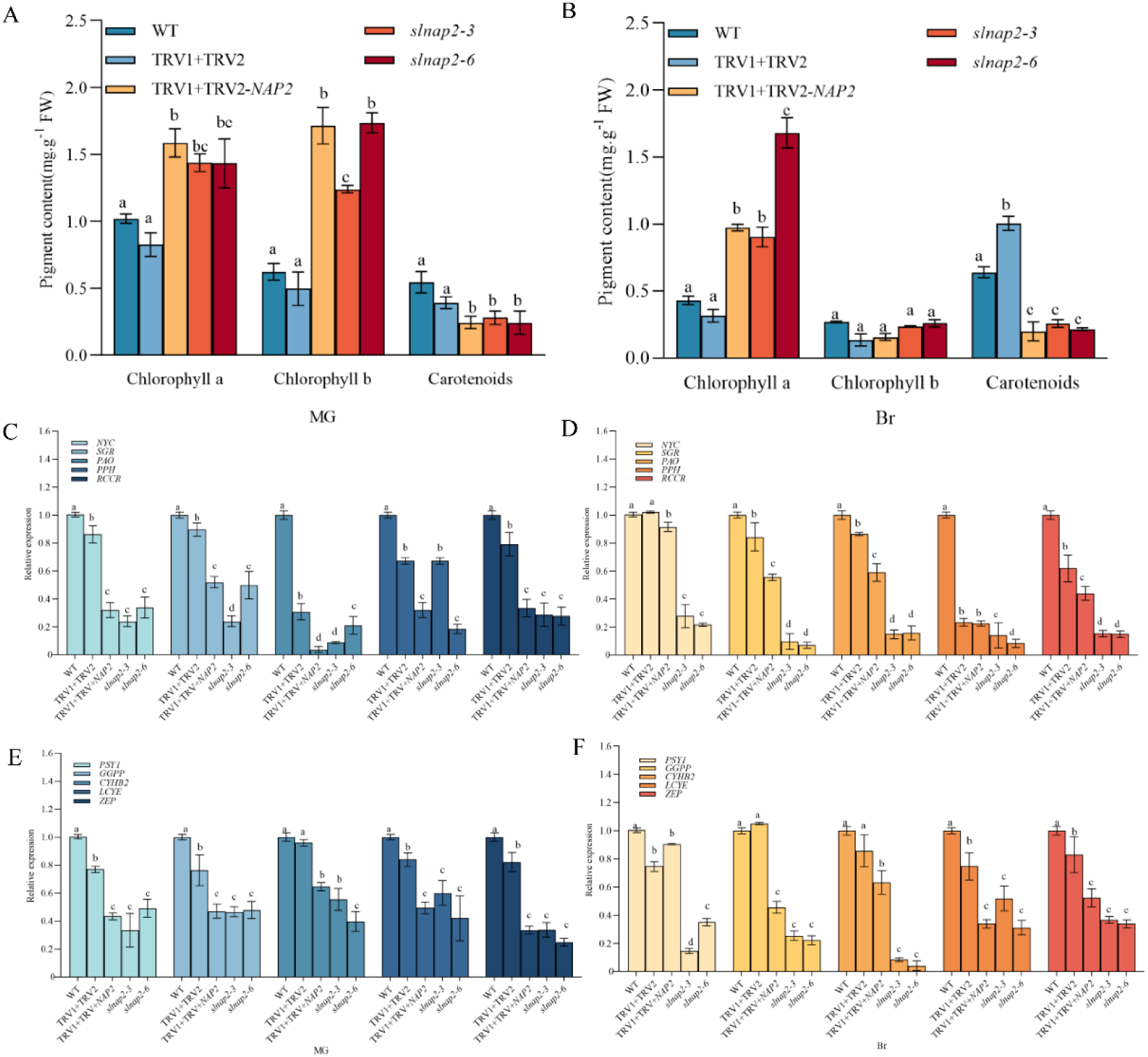
Pigment content and associated genes expression in the three lines at MG and Br stages. **A)** Chlorophyll a, Chlorophyll b and carotenoid content in the three lines at MG stage. **B)** Chlorophyll a, Chlorophyll b and carotenoid content in the three lines at Br stage. **C)** The transcript abundance of chlorophyll degradation-related genes in the three lines at MG stage. **D)** The transcript abundance of chlorophyll degradation-related genes in the three lines at Br stage. **E)** The transcript abundance of carotenoid synthesis-related genes in the three lines at MG stage. **F)** The transcript abundance of carotenoid synthesis-related genes in the three lines at Br stage. The different lowercase letters signify significant differences with a *p* <0.05, as evaluated by Duncan’s test.

### *SlNAP2* enhances the firmness in tomato fruit

To examine whether the *slnap2* mutants also influences fruit firmness to mediate ripening, we determined fruit firmness at the MG and Br stage in three lines. The results demonstrated that *SlNAP2* deletion mutant fruits were firmer than WT plants at MG stage, but there was no difference in the firmness of silenced plant fruits in comparison to WT fruits (Fig. 4A). Similarly, the firmness of silencing and mutant fruits was significantly more elevated at the Br stage than WT plants, and in particular, the firmness of *slnap2-3* and *slnap2-6* fruits was respectively increased by 88.2 %and 88.9 %. Subsequently, we examined the transcript levels of *PL8*, *MAN4*, *EXP*1, *TBG4* and *CEL2* in MG and Br stage fruit. Compared with WT plants, expression of *PL8*, *MAN4*, *EXP*1, *TBG4* and *CEL2* (cell wall degradation related genes) were sharply downregulated in *slnap2-3*, *slnap2-6* and TRV1+TRV2-*NAP2* at MG and Br stage (Fig. 4B). Interestingly, the *PL8*, *MAN4*, *EXP*1, *TBG4* and *CEL2* expression levels at the Br stage were downregulated by 90.1%, 92.4%, 85.6%, 92.7% and 98.2% in mutant fruits compared with WT fruits, respectively (Fig. 4C). Thus, the results demonstrate that the most striking feature of *slnap2*-regulated fruit ripening is the firmness at Br stage.

**Figure 4.**
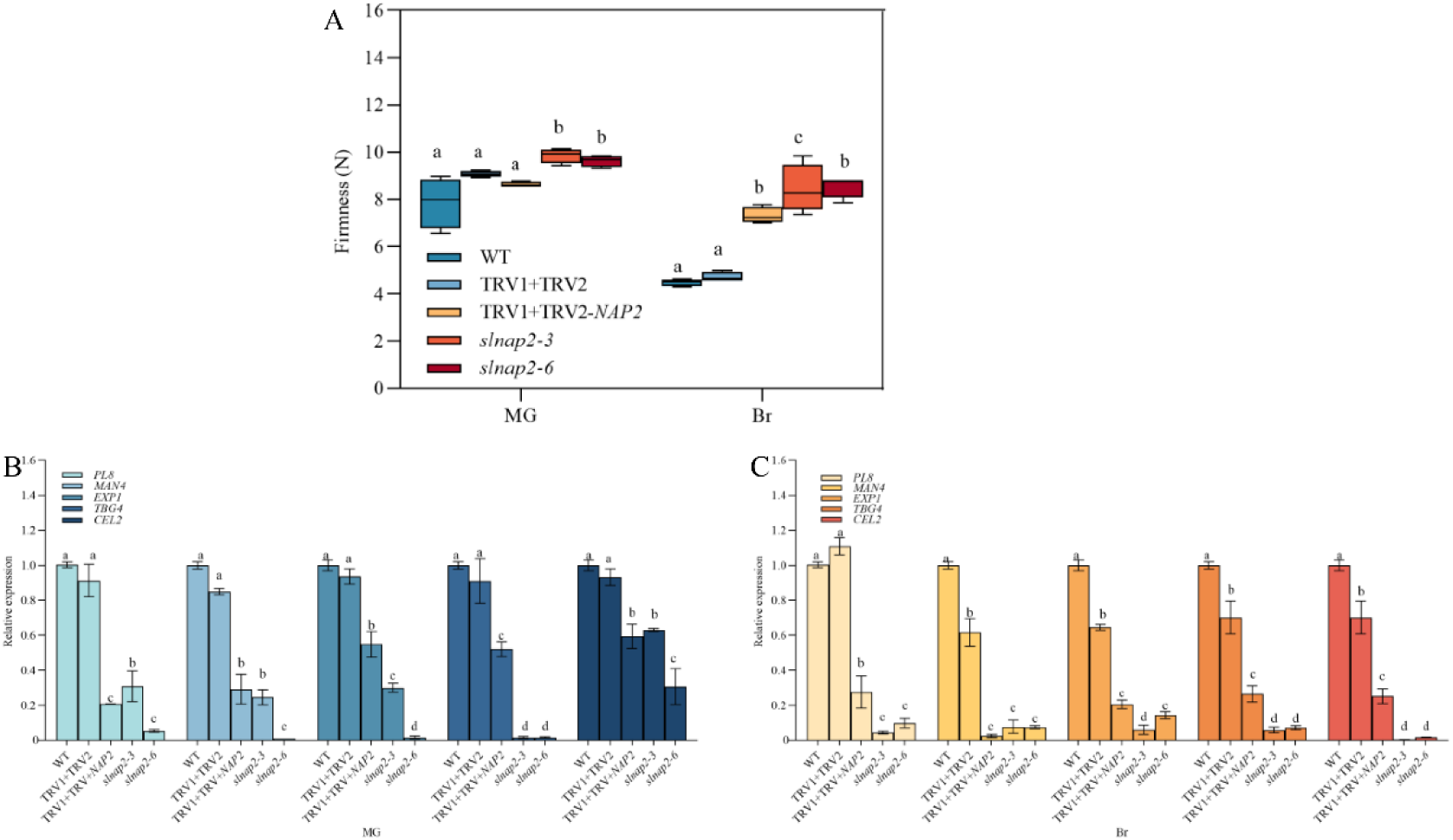
Fruit firmness and associated genes expression in the three lines at MG and Br stages. **A)** Fruit firmness in the three lines at MG and Br stages. **B)** The transcript level of cell wall metabolism-related genes in the three lines at MG stage. **C)** The transcript level of cell wall metabolism-related genes in the three lines at Br stage. The different lowercase letters signify significant differences with a * *p* <0.05, as evaluated by Duncan’s test.

### *SlNAP2* promotes the ethylene production in tomato fruit

To analyze exactly how fruit ripening is inhibited, the concentration of ethylene and ethylene production rate in *slnap2* mutant fruits were also investigated. There were no significant differences in ethylene concentration and ethylene production rate in fruit of the three lines in MG stage (Fig. 5, A and B). Notably, ethylene concentration and ethylene production rates were remarkablely more diminished in *slnap2-3*, *slnap2-6* and TRV1+TRV2-*NAP2* plants than in WT plants, especially at Br stage. Furthermore, expression of ethylene biosynthesis-associated genes (*ACS4*, *ACS2*, *ACO3* and *ACO1*) and ethylene signal transduction (*EIL3*) in fruits of mutant, silencing and WT plants was detected using qRT-PCR. Similarly, we observed that the expression of ethylene synthesis-related genes was dramatically decreased in mutant and silencing lines at MG and Br stage (Fig. 5, C and D). Moreover, the transcript abundance of *ACS2* in *slnap2-3* and *slnap2-6* mutants were down-regulated by 98.2% and 98.3% compared to the WT plants at Br stage (Fig. 5D). However, the transcript levels of *EIL3* in the *slnap2-3*, *slnap2-6* mutants and silencing lines, although significantly down-regulated, were not decreased to a large extent compared to ethylene synthesis-related genes at MG and Br stage. Therefore, we speculate that *EIL3* is crucial in *SlNAP2* mediating tomato fruit ripening in subsequent ethylene signaling. Additionally, to further confirm that *SlNAP2* can mediate fruit ripening, we further determined the transcript expression abundances of fruit ripening-related genes. The transcript abundances of *RIN*, *E4*, *NOR* and *E8* in *slnap2-3* and *slnap2-6* mutants were down-regulated in comparison to the WT plants at MG and Br stages (Supplemental Fig. S3). Especially at Br stage, the transcript expression abundances of ripening-associated genes in the *SlNAP2* knockout mutant were notably reduced compared to the WT. This further suggest that SlNAP2 is indeed participated in the process of tomato fruit ripening.

**Figure 5.**
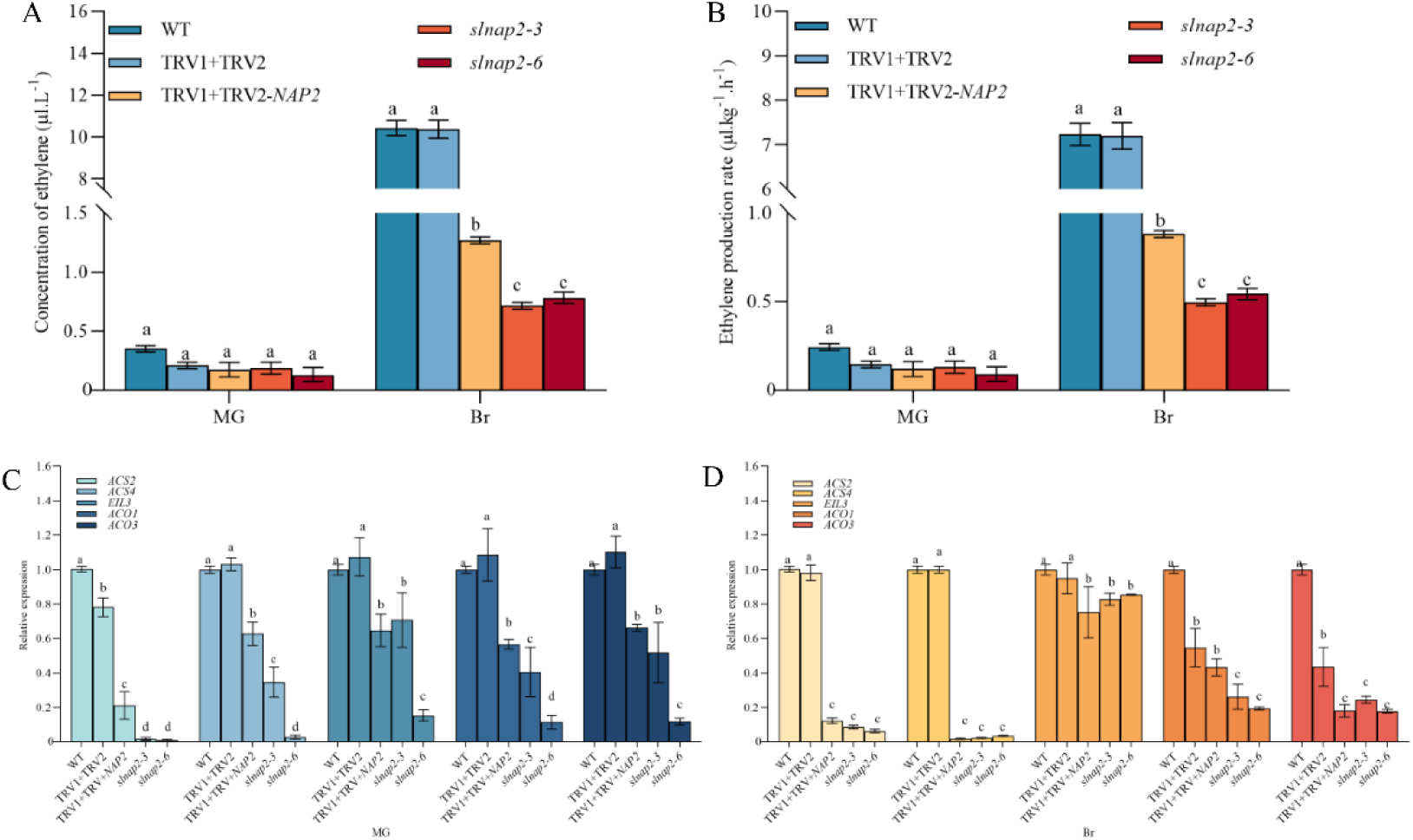
Ethylene production and associated genes expression in the three lines at MG and Br stages. **A)** Concentration of ethylene in the three lines at MG and Br stages. **B)** Ethylene production rate in the three lines at MG and Br stages. **C)** The relative expression level of ethylene synthesis-related genes in the three lines at MG stage. **D)** The relative expression level of ethylene synthesis-related genes in in the three lines at Br stage. The different lowercase letters signify significant differences with a * *p* <0.05, as evaluated by Duncan’s test.

### *SlNAP2* binds to the promoter of *SlACS2* to activate its transcriptional expression

To excavate the molecular process by which *SlNAP2* modulates fruit ripening, we utilized Y1H analysis. Combined with expression profiling of ethylene synthesis-related genes in *SlNAP2* knockout mutant fruits (Fig. 5C), we hypothesized that potential targets for downstream binding of SlNAP2 TFs are *SlACS2* and *SlACS1A*. Initially, we transformed the promoters of *SlACS2* and *SlACS1A* into yeast cells and suppressed the self-activation of the promoters using appropriate concentrations of AbA (Supplemental Fig. S4). Y1H analysis showed the promoter of *SlACS2* actually binds to SlNAP2 in *vitro* (Fig. 6A). However, *SlACS1A* failed to interact with NAP2, which suggests that *SlACS1A* is not a transcriptional binding target for SlNAP2 TFs (Fig. 6B). Subsequently, we further explored the gene expression of *SlACS2* and *SlACS1A* in *SlNAP2* knockout mutants and found that the transcript abundance of *SlACS1A* and *SlACS2* was dramatically reduced at both MG and Br stages (Fig. 6, C and D). We speculate that *SlNAP2* is mainly dependent on ethylene system II for the modulation of fruit ripening in tomato. To examine whether SlNAP2 directly influences the transcriptional activity of promoter of *SlACS2*, we obtained effector and reporter vectors for DLR assays (Fig. 6E). Compared with pGreen-62-SK-empty, pGreen-62-SK-SlNAP2 and pGreen-0800-LUC-*proACS2* co-transformed tobacco leaves with a higher relative luciferase activity (Fig. 6F). The results indicate that SlNAP2 bond to the promoter of *SlACS2* and activated at transcriptional level.

**Figure 6.**
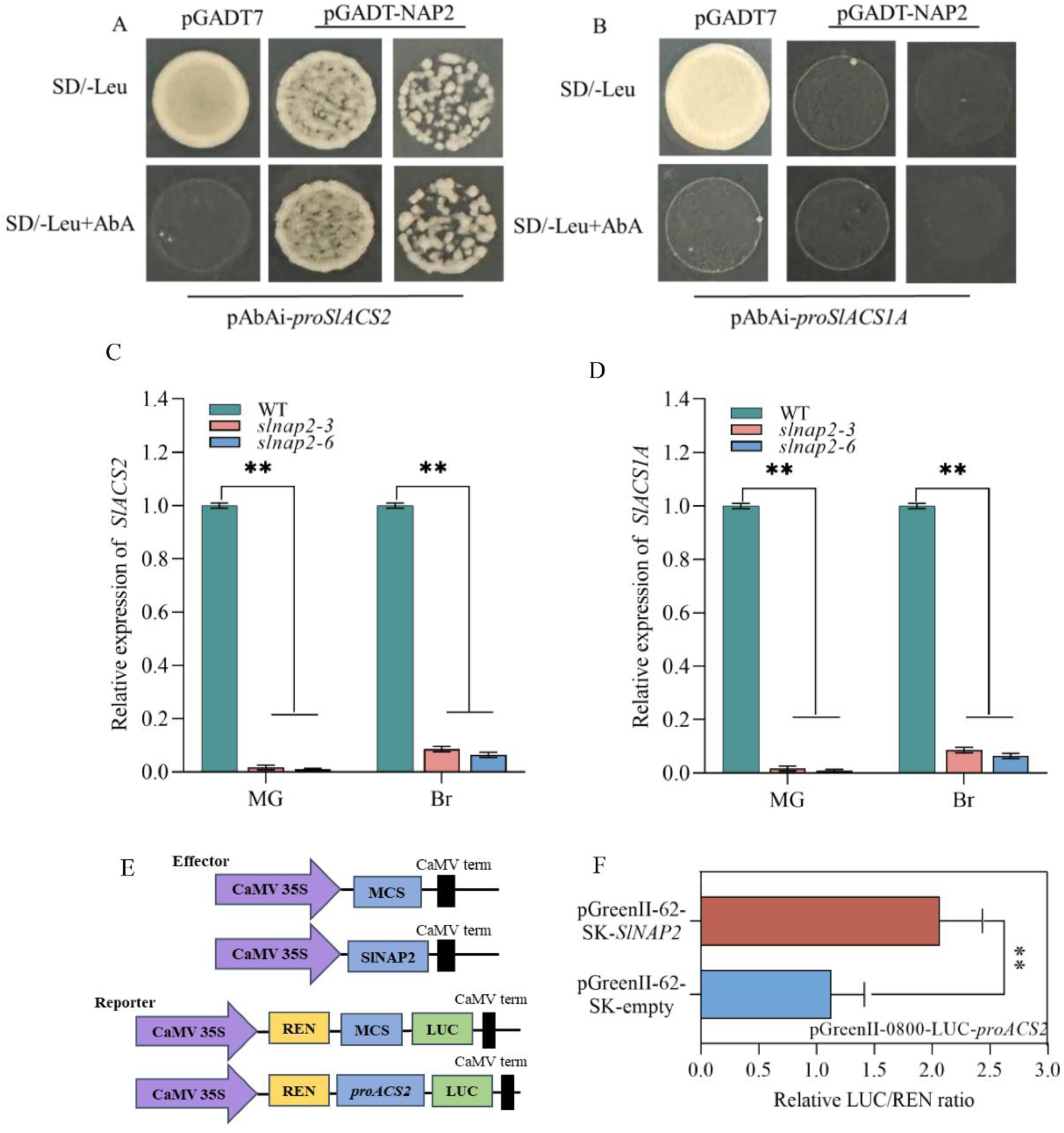
SlNAP2 binds to the *SlACS2* promoter. **A)** Y1H assay of SlNAP2 directly targeting the *SlACS2* promoter fragment. The SlNAP2 was linked to pGADT7 plasmid, and the *SlACS2* promoter were ligated to pBait-AbAi vector. **B)** Y1H analysis of SlNAP2 directly targeting the *SlACS1A* promoter fragment. The SlNAP2 was linked to pGADT7 vector, and the *SlACS1A* promoter were ligated to pBait-AbAi vector. **C)** The relative expression of *SlACS2* in WT and *slnap2* mutants at MG and Br stages. **D)** The relative expression of *SlACS1A* in WT and *slnap2* mutants at MG and Br stages. Asterisks indicate statistical significance evaluation by Student’s t-test, ** *p* <0.01. **E)** The CDS of SlNAP2 was cloned into the pGreenII-62-SK vector to generate the SlNAP2-62-SK effector. The CDS of SlNAP1 was cloned into the pGreenII 62-SK vector to generate the SlNAP1-62SK effector. The promoter of *SlACS2* was introduced into the pGreenII-0800-LUC vector to generate the proACS2-LUC reporter constructs. **F)** DLR assay system was used to measure the relative LUC/REN ratio. LUC, firefly luciferase activity; REN, Renilla luciferase. Asterisks indicate statistical significance evaluation by Student’s t-test, ** *p* <0.01.

### SlNAP2 interacts with SlEIL3 and increases the transcriptional activation of SlNAP2 on the promoters of SlACS2 gene

To additional determine how *SlNAP2* participates in ripening process, we employed the molecular biology techniques to explore its potential interaction proteins. Additionally, prior studies have shown a correlation between NAC transcription factors and ETHYLENE-INSENSITIVE-like protein (Shan et al., 2012). Considering the expression properties and patterns of the *SlNAP2* and *SlEIL3* genes in tomato fruit ripening. Therefore, we deduced that SlNAP2 could engage with SlEIL3 proteins. To verify this conjecture, the CDS of *SlNAP2* and *SlEIL3* were ligated into pGADT7 and pGBKT7 vectors for a Y2H assay (Fig. 7A). As SlNAP2 and SlEIL3 exhibited transactivation activity in yeast when linked with the pGBKT7 (Supplemental Fig. S5).

**Figure 7.**
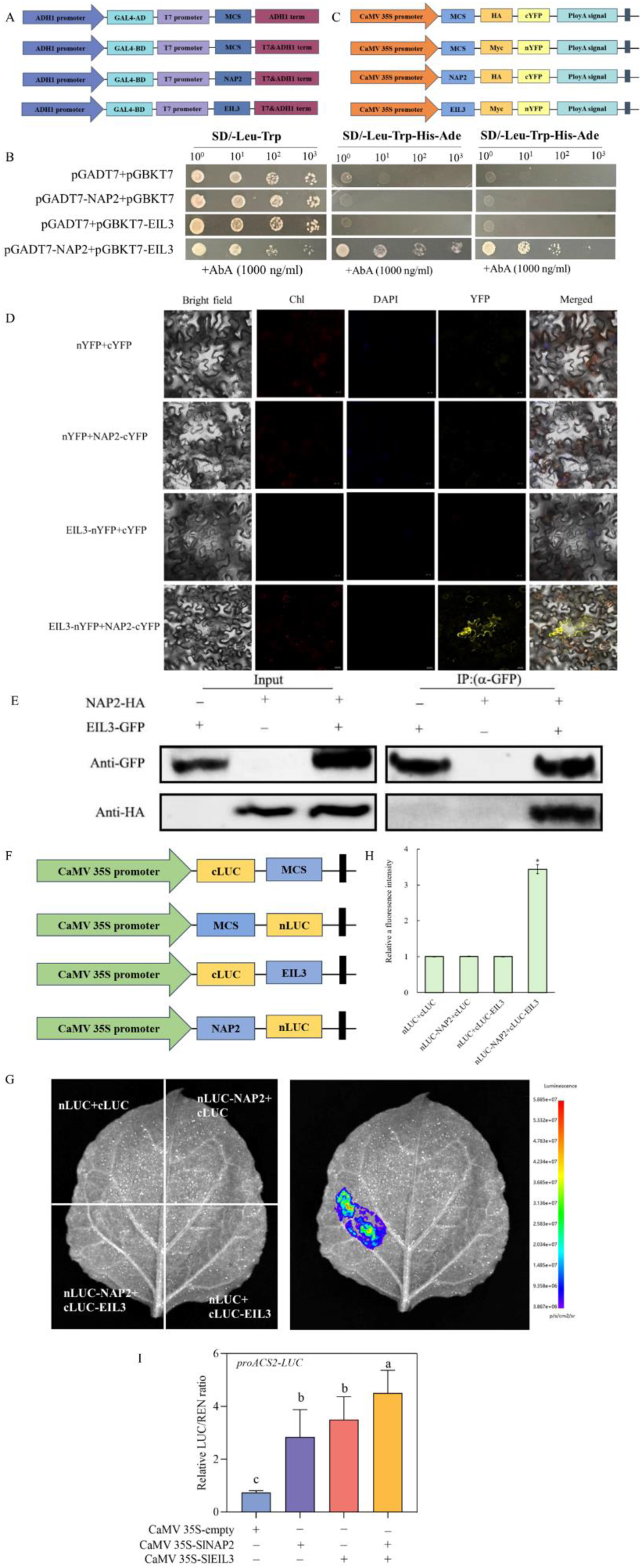
SlNAP1 interacts with SlEIL3. **A)** Schematic representation of the SlNAP2 and SlEIL3 ligated to pGADT7 and pGBKT7 vectors, respectively. **B)** Y2H analysis revealing the interaction between SlNAP2 and SlEIL3. Yeast cells co-transformed with pGADT7-NAP2+pGBKT7-EIL3 grew well. **C)** Schematic representation of the SlNAP2 and SlEIL3 fused to cYFP and nYFP bimolecular fluorescence complementary (BiFC) vectors, respectively. **D)** BiFC assays revealing the interactions between SlNAP2 and SlEIL3. **E)** Co-IP assay revealing SlNAP2 and SlEIL3 interaction. SlNAP2 and SlEIL3 were tagged with the HA and GFP tags. The precipitate was detected using anti-GFP and anti-HA antibodies. **F)** Schematic representation of the SlNAP2 and SlEIL3 fused to nLUC and cLUC luciferase complementation (LUC) vectors, respectively. **G)** Luciferase complimentary assay revealing the interaction of SlNAP2 and SlEIL3. **H)** The relative a fluorescence intensity was carried out using the PlantView software. Asterisks indicate statistical significance by Student’s t-test, * *p* <0.05. **I)** Transcriptional activity of SlNAP2 + SlEIL3 on promoter of *SlACS2*. Different lowercase letters indicate significant differences between groups. *p* < 0.05.

Subsequently, we simultaneously added AbA to the medium to suppress the transactivation activity of both vectors (pGADT7+pGBKT7-NAP2 and pGADT7+pGBKT7-EIL3) (Supplemental Fig. S5). Following, we performed Y2H experiments and found that yeast cells containing pGADT7-NAP2 and pGBKT7-EIL3 displayed excellent proliferation in the SD-Leu-Trp-His-Ade medium with AbA (1000ng/ml), confirming that SlNAP2 indeed interacted with SlEIL3 (Fig. 7B). To verify and validate the interaction between SlNAP2 and SlEIL3 detected in the Y2H assays, we conducted BiFC analysis using tobacco as a model system. SlNAP2 tagged with pSPYCE and SlEIL3 labeled to pSPYNE were co-infiltrated in tobacco leaves for transient expression (Fig. 7C). The YFP fluorescence was only observed in the cellular membrane of tobacco leaf cells with co-expressing NAP2-cYFP and EIL3-nYFP (Fig. 7D). Next, the CoIP assays by co-expressing EIL3-GFP and in tobacco leaves noted that NAP2-HA could be immunoprecipitated by EIL3-GFP in *vivo* (Fig. 7E). Meanwhile, we utilized a LUC imaging assay to explore the interaction between SlNAP2 and SlEIL3. SlNAP2-nLUC and cLUC-SlEIL3 constructs were co-injected into tobacco leaves (Fig. 7F). The bioluminescence was only observed in the tobacco leaf co-infiltrated SlNAP2 and SlEIL3 (Fig. 7G). Furthermore, the relative a fluorescence intensity was significantly elevated in comparison to the ratio achieved in the empty vector (Fig. 7H). These findings suggest that SlNAP2 physically interacted with SlEIL3. DLR assay demonstrated that the interaction of SlNAP2 with SlEIL3 enhanced the activity of the *SlACS2* promoter compared to CaMV 35S-SlNAP2 alone (Fig. 7I). Altogether, SlEIL3 may cooperate with SlNAP2, and further increase the binding activity of SlNAP2 on the promoters of *SlACS2* gene.

## Discussion

### *SlNAP2* advances the ripening of tomato fruit

In recent, the functional and structural properties of NAC TFs have been studied and elucidated (Diao et al., 2020). NAC TFs, which comprise ATAF, NAM, and CUC members, form a plant-specific superfamily (Li et al., 2024). The tomato genome includes 101 NAC TFs (Ma et al., 2018). The different members of NAC TFs are participated in different physiological processes and perform a variety of functions in plants, containing plant growth and development and response to environmental stress (Olsen et al., 2005). In the current study, WT fruits entered the Re stage while *SlNAP2* mutant fruits were still stagnant in the Br stage (Fig. 2), indicating that knockout of *SlNAP2* obviously inhibited fruit ripening in tomato. Currently, there are abundant research to explore the regulatory roles of NAC TFs in tomato ripening. Scientists are investigating the signaling network that regulates fruit ripening, including upstream TFs and downstream target genes, as well as downstream abundant signaling proteins. In this study, we have identified that SlNAP2 accelerated tomato fruit ripening, acted directly on the promoter region of downstream target genes (*SlACS2*), and physically interacted with EIL3, a protein associated with ethylene signal transduction (Fig. 8).

**Figure 8.**
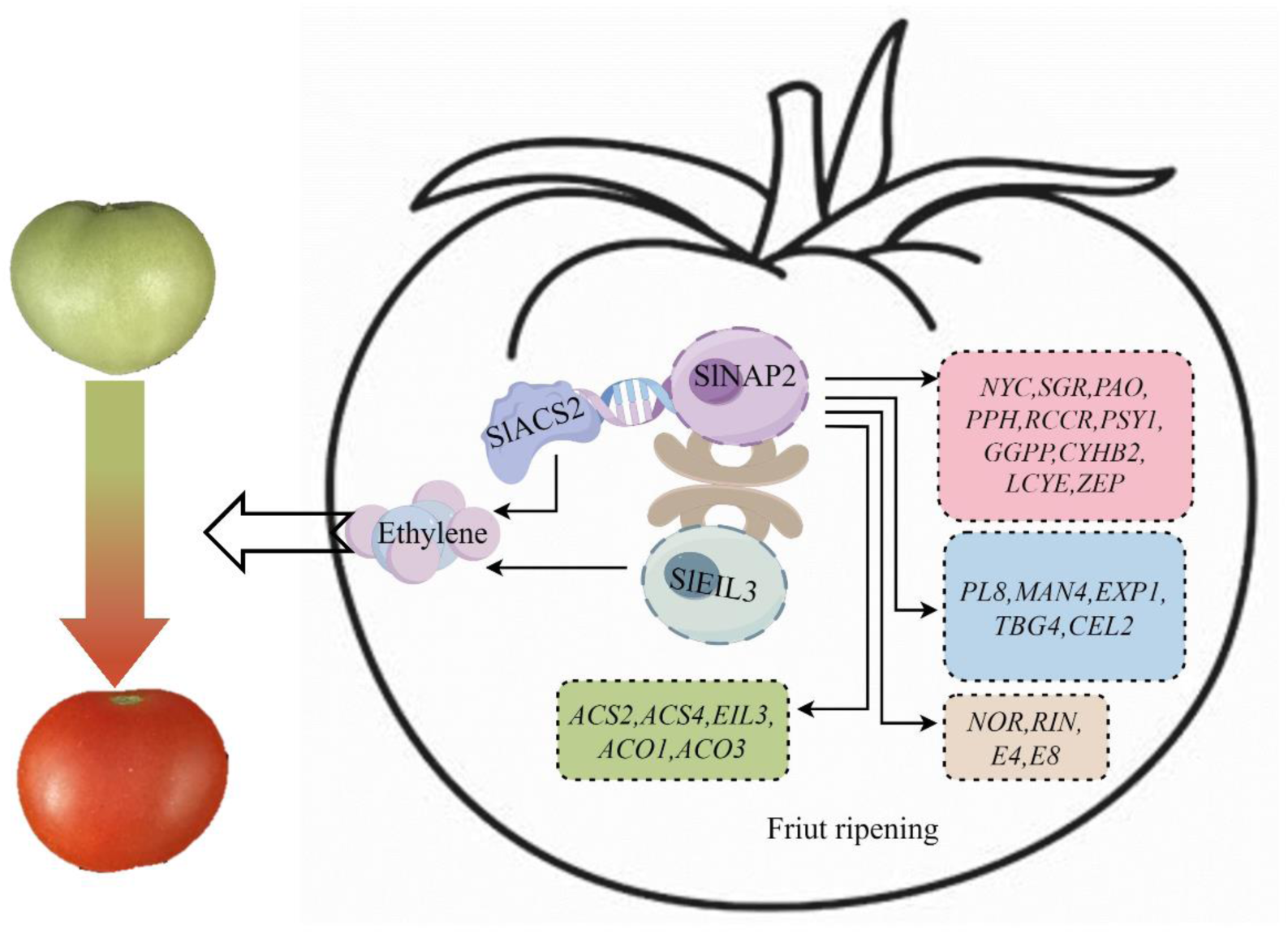
A schematic of the working model for the modulation of tomato fruit ripening by the SlNAP2 transcription factor. SlNAP2, a NAC transcription factor, directly binds to *SlACS2* to activate its expression and promote ethylene production, which in turn interacts with EIL3 to enhance the function of ethylene in tomato fruit ripening. Transcript levels of genes associated with chlorophyll degradation, carotenoid synthesis and cell wall metabolism are also differentially affected in *slnap2* knockout mutants. However, whether SlNAP2 also directly regulates these genes is currently unknown.

This lays the foundation for constructing the tomato ripening network and provides new evidence that NAC TFs and ethylene synergistically regulate tomato fruit ripening.

Besides the classical NOR, a number of NAC family genes have been shown to induce fruit ripening via transcriptional regulation (Kou et al., 2016; Gao et al., 2022; Feng et al., 2023). Gao et al. (2022) generated CR-*nor-like1*mutants in tomato utilizing CRISPR/Cas9 gene editing technology and observed significant inhibition of ripening. They found that the mutant fruits were unsuccessful in turn completely red, remaining orange-red at the final stage of ripening. Ripening defects were observed in the pericarp of tomato plants in the *SlNAC4* RNAi lines, indicating that the suppression of *SlNAC4* gene expression affected normal fruit development and maturation processes. (Zhu et al., 2014). Similarly, the flowering process was delayed and the transition to color break was significantly inhibited in the *SNAC9* KO line, which in turn substantially hindered the fruit ripening process (Feng et al., 2023). In this study, we observed that as WT fruits entered the Re stage, fruits of the *SlNAP2* silencing lines and the mutants remained stagnant in the Br stage (Fig. 2). These findings clearly show that NAC TFs are participated in the process of tomato fruit ripening. Specifically, NAP2 has been identified as a positive transcription activator within the ripening process. This underscores the importance of NAC TFs in the modulation and coordination of key developmental stages in tomato.

### *SlNAP2* influences chlorophyll and carotenoid metabolism in tomato fruit

The series of color transformation from green to orange to red during tomato fruit ripening are directly affected by pigment development in fruit (Klee and Giovannoni, 2011). The content of chlorophyll, carotenoids and lycopene is one of the distinctive markers of ripeness of tomato fruits. In our study, the *slnap2-3* and *slnap2-6* mutant fruits had marked elevation in chlorophyll content at MG stage compared to WT plants (Fig. 3). Meanwhile, carotenoid content was obviously lower in the *slnap2-3* and *slanp2-6* mutants than in the WT plants both at MG and Br stages (Fig. 3). Consistent with our data, Gao et al. (2022) also explored that total chlorophyll content was significantly augmented, and carotenoid and lycopene content was significantly reduced in *nor-like1* mutant fruits. Similarly, the SlNAC4 RNAi lines exhibited remarkably higher total chlorophyll content and obviously lower total carotenoid content compared to the WT (Zhu et al., 2014). Feng et al. (2023) also indicated that the total chlorophyll content and carotenoid content in CRISPR-SNAC9 tomato fruits were higher and lower than those in WT fruits, respectively. These results deduce that NAC TFs play a significant and crucial role in mediating pigment content, which in turn influences the fruit ripening process. By modulating the synthesis and accumulation of pigments, NAC TFs contribute to the color development that is crucial for the ripening and overall appearance of the fruit. This suggests a vital link between the action of these transcription factors and the visual and physiological changes that occur during fruit maturation. The expression of *PSY1* is boosted during the ripening process, thereby promoting the generation of carotenoids that contribute to fruit pigmentation (Kachanovsky et al., 2012). Previous studies have found that unlike other plants such as Arabidopsis, rice or pepper, which rely on a single GGPPS isoform for the production of the required plastid-like isoprenoids, the production of a majority of the GGPP in the chloroplasts of tomato is dependent on two GGPPS homologues (Zhou et al., 2017; Wang et al., 2018; Barja et al., 2021). Their experiments also confirmed that *PSY1* expression was significantly increased in fruit ripening. However, when we knocked out *SlNAP2* gene, *PSY1*, *GGPP*, *CYHB2*, *LYCE*, and *ZEP* expression level were remarkably reduced. Furthermore, RNA-seq data showed significant down-regulation of *SlGgpps2* and *SlSGR1* genes participated in carotenoid accumulation and chlorophyll degradation in the *nor-like1* knockout mutant (Gao et al., 2022). They also demonstrated that NOR-like1 directly controlled the target gene (*SlSGR1* and *SlGgpps2*) initiation, thereby influencing ripeness of tomato fruit. However, our study has not yet demonstrated that SlNAP2 can directly activate or suppress pigment metabolism-related genes to mediate fruit ripening.

### *SlNAP2* affects the firmness in tomato fruit

The firmness of climacteric fruit as a favourite horticultural product for consumers is also an extremely important quality index. Fruit ripening is often accompanied by flesh softening, a process that is mediated by a range of genes engaged in cell wall metabolism through remodelling, degradation and polymerisation of cell wall components (Liu et al., 2021). Previous study demonstrated that CRISPR/Cas9 technology-mediated *slpg* mutant fruits were less firmness and had higher physiological loss of water than WT fruits (Nie et al., 2022). This suggests that knockout of the polygalacturonase (*PG*) gene could indeed alter fruit firmness, in contrast to our experiments where knockout of the *SlNAP2* gene resulted in an enhancement in fruit firmness (Fig. 3). Gao et al. (2022) showed that the firmness of mutant fruits increased following knockout of *nor-like1* gene, which is consistent with our finding. In parallel, they found that the mutant cell walls were darker in color and the microfilaments were more tightly arranged. Furthermore, overexpression of *SlNAC6* expedited tomato fruit ripening, and an early initiation of *PG*, *CEL2* and *EXP1* expression was found in the SlNACC6-OE lines (Jian et al., 2021); therefore, they hypothesized that NAC6 could promote tomato ripening by mediating the untimely transcription of *PG*, *CEL2* and *EXP1*. In line with this, CRISPR/Cas9-mediated *SlNAP2* mutant also enhanced the fruit firmness, and the *PL8*, *MAN4*, *EXP*1, *TBG4* and *CEL2* expression levels were downregulated in mutant fruits at Br stage (Fig. 4). Meanwhile, the relative expression levels of *SlPG2a*, *SlPL*, *SlCEL2*, and *SlEXP1* were apparently decreased by at least 89% in *nor-like1* deletion mutant (Gao et al., 2022). Feng et al. (2023) also found that CRISPR/Cas9-mediated SNAC9 mutants had high fruit firmness. In this study, expression of *PL8*, *MAN4*, *EXP1*, *TBG4* and *CEL2* was sharply downregulated in *slnap2-3*, *slnap2-6* and TRV1+TRV2-*NAP2* at MG and Br stage (Fig. 4). Similarly, strand specific mRNA of *slnam* mutant and OE-SlNAM lines fruit revealed the enrichment of *EXP1* and *CEL2* in the differential genes, and RT-PCR revealed that the transcript abundance of *EXP1* and *CEL2* was also significantly reduced in the *slnam* mutant. In addition, a NAC TFs, NOR-Like targeted the *SlPG2a*, *SlPL*, *SlCEL2* and *SlEXP1* promoters to modulate cell wall metabolism in ripening of tomato fruit (Gao et al., 2022). Therefore, NAC TFs are potentially participated in the modulation of cell wall metabolism. By influencing the biochemical pathways that modify the cell wall structure, NAC TFs help to control the softening of the fruit, which is a key characteristic of ripening.

### SlNAP2 interacts with *ACS2* and *EIL3* to regulate tomato fruit ripening

The initiation of ethylene generation is crucial for the climacteric fruit ripening (Alexander, 2002). Previous studies revealed that *SlACS6* and *SlACS1A* were primarily participated in system I ethylene, adversely influenced by ethylene itself, whereas system II ethylene depended on *SlACS2*, *SlACS1A*, and *SlACS4*, which were notably stimulated by ethylene from MG stage onward (Barry et al., 2000; Liu et al., 2015). Furthermore, ethylene perception and the following signaling pathways are well-documented in Arabidopsis (Binder, 2020). ETHYLENE INSENSITIVE2 (EIN2) protein is central in the ethylene signaling cascade, as EIN2 deficiency in Arabidopsis blocks all ethylene responses (Alonso et al., 1999). Moreover, the EIN2 (C-terminal region) is translocated from the endoplasmic reticulum to the nucleus to stimulate the EIL1 and EIN3 (Ju et al., 2012; Wen et al., 2012; Qiao et al., 2012). In Arabidopsis, the TFs EIN3 and EIL1 play crucial roles in modulating the plant’s response to ethylene, acting as central components in the ethylene signaling network (Chao et al., 1997). Previous study found that *ACS2* and *ACS4* were expressed at high levels in tomato fruit, and that co-deletion of *ACS4* and *ACS2* blocked the generation of the ethylene burst essential for the ripening of tomato fruit (Hoogstrate et al., 2014). Analogously, our findings demonstrated that the *SlACS2* and *SlACS4* were expressed at low levels in NAP2-deficient tomato fruits (Fig. 5). Meanwhile, TFs NAP2 could directly ligate to the *SlACS2* promoter to stimulate its transcriptional activity and thus its gene expression. Furthermore, further exploration revealed that NAP2 could interact with EIL3 and increase the transcriptional activation of SlNAP2 on the promoters of *SlACS2* gene (Fig. 7). We hypothesize that NAP2 might modulate the model of fruit ripening: NAP2 could directly influence the *ACS2* promoter, promote its transcriptional expression, and further interact with EIL3 to form a complex to promote tomato fruit ripening (Fig. 8). Gao et al. (2022) revealed that NAC TFs, NOR-like1 directly interacted with the *SlACS4* and *SlACS2* promoters to modulate ethylene generation. Intriguingly, methyl jasmonate accelerated ethylene synthesis in kiwifruit by preferentially activating *AdACS1* and *AdACS2* via inducing *NAC2/3* gene (Wu et al., 2020). Recently, NAC TFs in tomato, SlNAM1, positively influenced ethylene biosynthesis by directly ligating to two key ethylene biosynthesis genes: *SlACS4* and *SlACS2* (Gao et al., 2021). Taken together, NAC TFs facilitate fruit ripening by mediating ethylene synthesis, which might be mainly dependent on affecting ethylene system II. Shan et al., (2012) demonstrated that the NAC TFs in banana, MaNAC1/2, intimately interacted with the ethylene signaling component, the EIL3 protein, EIL5, which is downregulated during ripening. This suggests that NAC TFs might simultaneously be able to interact with ethylene signaling components involved in fruit ripening. Similarly, we also found that SlNAP2 could interact with EIL3. This observation supports further investigation into how NAC TFs and components of the ethylene signaling network collaboratively regulate the ripening process of tomato fruits. Understanding the interaction between these elements could reveal more about the complex regulatory mechanisms that control fruit maturation and potentially lead to strategies for improving fruit quality and shelf life. Through a combination of the phenotypic and molecular assay results described above, we have proposed a possible working hypothesis for the role of NAC TFs, SlNAP2 and ethylene in mediating fruit ripening (Fig. 8). SlNAP2 mediates tomato fruit ripening by binding to *SlACS2* to activate its expression and then interacting with downstream EIL3 to influence ethylene biosynthesis and ethylene signaling effects. The transcript levels of genes associated with carotenoid and chlorophyll metabolism and cell wall metabolism are also potentially affected by the *SlNAP2* deletion; however, it remained unclear whether these genes are also directly regulated by SlNAP2. Our data contribute to understanding the regulatory role of NAC TFs in ethylene during tomato fruit ripening, thereby enhancing our knowledge of the ripening regulatory network that governs tomato fruit maturation.

## Materials and methods

### Plant materials and growth conditions

Tomato (*Solanum lycopersicum* cv. ‘Micro-Tom’) cultured in a growth chamber was regarded as the wild type (WT). The WT and knockout mutants were cultivated at 22/20°C with 80% relative humidity for 16/8 h (day/night) under a luminous intensity of 400 μmol m^-2^ sec^-1^. To accurately measure ripening time, anthesis flowers of WT, mutant and transiently silenced plants were marked and the duration from anthesis to fruit ripening was noted until the color transition occurred. Fruit ripening stages were categorized as mature green (MG), breaker (Br), 5 days after breaker (Br+5), and red ripening (Re) stages. In addition, tobacco (*Nicotiana benthamiana*) used for transient expression was cultivated at 25°C, 16/8 h (day/night) in a growth room. All freshly collected fruit samples were stored at -80°C until necessary to remove them for use.

### CRISPR/Cas9 gene editing

Tomato *slnap2* mutant material was obtained using CRISPR/Cas9 gene editing and tissue culture techniques. The 20 bases upstream of NGG in the *SlNAP2* gene coding sequence (CDS) region was selected as the target sequence on the CRISPR-P website (http://cbi.hzau.edu.cn/cgi-bin/CRISPR). The target sequence was reverse complemented to sgRNA and imported into CRISPR/sgRNA vectors, and then ligated into the CRISPR/Cas9 binary vector pHEE401. The construct was introduced into tomato cv. ‘Micro-Tom’ by infecting leaf explants with *Agrobacterium tumefaciens* GV3101. Based on their hygromycin B resistance, transgenic tomato lines were screened. After successful transformation, leaf DNA was extracted using Fast Plant DNAzol (Zhenbai Biotechnology, Hangzhou, China). DNA fragments (500bp) containing upstream and downstream regions of the target sequence were amplified. The PCR reaction solution was sequenced by a sequencing company, and after sequence alignment, screened for two pure and stable genetic lines. The results of off-target site detection are provided in Supplemental Fig. S2.

### Virus-induced gene silencing

To obtain constructs for virus-induced gene silencing (VIGS), a 200-300 bp cDNA sequence of SlNAP2 was inserted into the pTRV2 vector using gene-specific primers (Table S1) and then heat-stimulated for transfer into *Agrobacterium tumefaciens* strain GV3011. Tomato fruits were injectively infiltrated with *Agrobacterium* carrying the TRV construct as outlined by Yuan et al. (2016). Additionally, *Agrobacterium tumefaciens* GV3101 carrying TRV-VIGS vector was growed in Luria-Bertani (LB) medium supplemented with MES (10 mM) and acetosyringone (200 µM) at 28°C with suitable antibiotics. After 16 h of incubation with shaking, the bacterial fluid was collected, resuspended in infiltration buffer (10 mM MgCl_2_, 10 mM MES, pH 5.6, 200 µM acetosyringone) to achieve a final OD600 of 0.8-1 (pTRV1 or pTRV2 and pTRV2-SlNAP2), and left to rest for 3 h at 28 °C before infiltration. A mixture of resuspended pTRV1 and pTRV2 or pTRV2-SlNAP2 in a 1:1 ratio was introduced into the pericarp of tomato fruits using a 1ml syringe. Subsequently, we detected the expression of *SlNAP2* in the fruit three days after infiltration by qRT-PCR.

### Total RNA isolation and quantitative real-time PCR (qRT-PCR) analysis

Total RNA was isolated from tomato fruits (MG and Br) using TRIzol reagent (Invitrogen, Shanghai, China) following the guidance instructions. Briefly, the fruit samples were pulverized to micronize using liquid nitrogen in a pre-cooled mortar and pestle. Subsequently, TRIzol (1 mL) was poured in the mixture and incubated at 4°C for 10 min. Chloroform (200 µL) was introduced into the mixture, followed by incubation at 4°C for 5 min. The supernatant was obtained by centrifuging at 12000 *g* for 15 min at 4°C. Afterward, an equal volume of isopropanol (about 600µL) was added to the mixture and then allowed to stand at -20°C for more than 1 h. The mixture was transferred to an adsorption column centrifuged at 12000 *g* for 15 min at 4°C. The fruit RNA was rinsed twice with 75% ethanol and ultimately dissolved in RNase-Free ddH_2_O.

The reverse transcription was performed using *Evo M-MLV* RT premix (Accurate Biology, Changsha, China). The qRT-PCR analysis utilized primers listed in Table S1 and was conducted in the Light Cycler™ 96 Real-Time PCR System according to the manufacturer’s guidelines with the SYBR Green Premix Pro Taq HS qPCR Kit (Accurate Biology, Changsha, China). The tomato actin gene (Solyc03g078400) was served as an internal control for determining relative expression levels. Gene relative expression values were determined using the 2^-ΔΔCt^ method (Livak and Schmittgen, 2001). Each sample was analyzed with three biological replicates.

### Subcellular localization

The *SlNAP2* gene open reading frame (ORF) was inserted into the *XbaI* and *KpnI* sites in the pSuper1300 vector. The SlNAP2-GFP fusion expression vector was constructed using a super-promoter for regulation. The SlNAP2-GFP recombinant vectors were transiently introduced into tobacco (*N*. *benthamiana*) leaves using GV3101 strain. Dark incubation for 24 h followed by normal incubation for 72 h, GFP fluorescence in tobacco leaves was detected and captured with a confocal microscope (Zeiss LSM 710, Leica Wetzlar, Germany). The primers utilized in this study are detailed in Supplementary Table S1.

### Transcription activation analysis

The *SlNAP2* was linked to the pGBK-BD vector to generate the pGBK-NAP2 construct. Subsequently, pGBK-empty and pGBK-NAP2 construct was introduced into yeast strain Y2H. Well-grown transformant was picked on synthetic dropout (SD)/- tryptophan (Trp) deletion medium. The transformants were propagated in YPDA medium and diluted to OD600 = 0.2 with 0.9% NaCl (w/v) and then spotted on SD/- Trp/-adenine (Ade)/-histidine (His) deletion medium according to the concentration gradient of the dilution. The yeasts were grown at 30°C. Details about the primers can be obtained in Supplementary Table S1.

### Chromaticity analysis of tomato fruit

A chroma meter CR-400 (Konica Minolta Inc, Japan) was utilized to evaluate the tomato fruit color. Before detection, calibration with a standard whiteboard was performed prior to measuring the L, a, and b values of the WT, mutant and silenced fruit. L*, a* and b* were recorded for each fruit at three independent points.

### Measurement of pigment content

The content of total chloroplast pigments was determined with slight modifications with reference to Yang et al. (2020). The total chloroplast pigments were extracted using 95% ethanol (10 mL) in darkness for 24 h. The absorbances were determined at 665 and 649 nm by UV spectrophotometer (Shimadzu UV-1780, Kyoto, Japan). The pigment content was computed using the following formula: chlorophyll a = 13.95 × A_665_ – 6.68 × A_649_; chlorophyll b = 24.96 × A_649_ – 7.32 × A_665_. Carotenoids were extracted in darkness with 80% acetone for 24 h. The absorbance was determined at 470 nm. The carotenoid content was calculated using the provided formula: carotenoid = (1000 × A_470_ – 2.05× chlorophyll a – 114.8 × chlorophyll b)/245.

### Measurement of firmness

Tomato fruit firmness was measured with a Universal TA texture analyzer (Tengba Inc, Shanghai, China), following the guidelines provided by the manufacturer. Each assay involved a minimum of nine biological replicates, with each replicate obtained through independent sampling. The parameters were taken in Newtons (N).

### Measurement of ethylene content and ethylene production rate

To determine ethylene levels, fruits from the three lines were collected at MG, Br stages and allowed to place at room temperature for 2 h to avoid recording the transient “wound ethylene” that is typically emitted shortly after harvesting. The fruits were enclosed in 500 mL gas-tight bottles and left at room temperature for 12 h. Gas (1 mL) sample was injected and analysed by SP-3420 gas chromatograph (Beifenruili, Beijing, China). The ethylene concentration was assessed by comparing the peak area of the sample with that of a known concentration ethylene standard. The ethylene production rate represents the quantity of ethylene generated per gram of tomato fruit per hour in a confined space. Each measurement was conducted using a minimum of three replicates.

### Yeast one-hybrid assays

The *SlNAP2* full fragment was isolated from tomato cDNA and ligated into the pGADT7 with *EcoRI* and *BamHI* digestion. The promoters of *SlACS1A* and *SlACS2* were inserted into the vector via *KpnI* and *XhoI* digestion. The resulting pBait-AbAi-promoter vector was then linearized using *BbsI* and introduced into the yeast Y1H strain. Subsequently, the positive plaques were selected on SD/- uracil (Ura). Suitable concentrations of aureobasidin A (AbA) were picked to suppress self-activation. The pGADT7-NAP2 plasmid was then transformed into a positive pBait-AbAi-promoter yeast strain, spotted on SD/- leucine (Leu) medium (+/-AbA), and diluted in a concentration gradient, and cultured for 3 d. Primers details are available Supplementary Table S1.

### Transient dual-luciferase expression (DLR) assays

The promoters of *SlACS2* were inserted into the pGreenII-0800-LUC reporter vector. The CDS of SlNAP2 was cloned into the pGreenII-62-SK effector vector. Subsequently, the reporter vector and effector vector were co-infiltrated into tobacco leaves. After 72 h, the DLR assay was measured using a dual luciferase assay kit (Promega, WI, USA).

### Yeast two-hybrid assays

The full fragment of *SlNAP2* and *SlEIL3* were inserted into the pGBKT7 vector via the digestion of *EcoRI* and *BamHI*. Subsequently, the recombinant plasmid and pGADT7 plasmid using the PEG/LiAC approach were co-introduced into the Y2H yeast strain, and the appropriate concentration of AbA was screened to inhibit the self-activation on SD/-Leu/-Trp/-His/-Ade medium. The CDSs of *SlNAP2* and *SlEIL3* were inserted into pGADT7 and pGBKT7 vectors, respectively, to generate different baits and preys. Subsequently, the bait vector and prey vector were co-transfected into the yeast strain Y2H. Yeast was cultured on SD/-Leu/-Trp-deficient medium, and interactions were explored by spot-plating on SD/-Leu/-Trp/-His/-Ade medium containing the appropriate concentration of AbA according to the concentration gradient of the dilution. The primers utilized are applied in Supplementary Table S1.

### Bimolecular fluorescence complementation assay

The bimolecular fluorescence complementation (BiFC) assay was employed to observe interactions. The CDS of *SlNAP2*, lacking a stop codon, was integrated into the pSPYCE-35S vector to produce the NAP2-cYFP construct. Then, the termination codon-free *SlEIL3* was inserted into the pSPYNE-35S plasmid to create the EIL3-nYFP construct. These recombinant plasmids of NAP2-cYFP and EIL3-nYFP constructs were then introduced into GV3101 strain and subsequently injected in tobacco leaves. After dark cultivation for 96 h, YFP fluorescence was viewed under a confocal microscope at 4 d (Zeiss LSM 710, Leica Wetzlar, Germany). Details on the primers are provided in Supplementary Table S1.

### Luciferase complementation assay

The luciferase complementation assay (LUC) assay was used for detecting protein interactions *in vivo*. The CDS of *SlNAP2*, excluding the stop codon, was ligated into the pCAMBIA1300-nLUC plasmid to create the NAP2-nLUC construct; similarly, SlEIL3 gene without stop codon was inserted into the pCAMBIA1300-cLUC plasmid to create the EIL3-cLUC construct. The engineered plasmid was introduced into *Agrobacterium tumefaciens* GV3101 and subsequently infiltrated into tobacco leaves. Following 96 h of cultivation, the LUC signal in tobacco leaves was viewed by an PlantView100 at 4 d (BLT photon technology, Guangzhou, China) *in vivo*. The primers utilized in this assay are detailed in Supplementary Table S1.

### Coimmunoprecipitation analysis

The CDSs of *SlNAP2* and *SlEIL3* without termination codon were cloned into pAC004-HA and pSuper1300-GFP vectors to generate NAP2-HA and EIL3-GFP constructs, respectively. *Agrobacterium* containing NAP2-HA and EIL3-GFP were cotransformed into tobacco leaves. Co-IP was conducted following previously established methods (Ji et al., 2021). Briefly, following a 96 h incubation period, total proteins were isolated from tobacco leaves and subsequently bound to GFP agarose beads (ChromoTek, Wuhan, China). Input proteins and their corresponding immunoprecipitates were identified through the process of immunoblotting, utilizing specific antibodies against GFP or HA to detect the presence and interaction of these proteins. The primers utilized in this experiment are detailed in Supplementary Table S1.

### Statistical analysis

Data was statistically analyzed using SPSS software, Excel, GraphPad Prism. Pairwise comparisons were made using the Student’s t-test (*p* < 0.05), while multiple comparisons were made using one-way ANOVA and Duncan’s multiple polarity test.

## Supplemental data

The following materials related to this article are obtained in the online version.

**Supplemental Figure S1.** The analysis of subcellular localization of SlNAP2. A, Phylogenetic analysis of SlNAC1, SlNAM, AtNAP, SlNAP1 and SlNAP2 protein. B, Schematic representation of SlNAP2 and GFP fluorescence vectors. C, The subcellular localization of SlNAP2 in tobacco. D, Transcriptional activation of SlNAP2.

**Supplemental Figure S2.** Sequence identification of stable transgenic material after *SlNAP2* knockout.

**Supplemental Figure S3.** Fruit ripening-related genes expression in WT, silencing lines, and *SlNAP2* mutants. The different lowercase letters signify significant differences with a * *p* <0.05, as determined by Duncan’s test.

**Supplemental Figure S4.** The screen of optimal AbA concentration to inhibit the self-activation of *SlACS2* and *SlACS1A* promoters. AbA: Aureobasidin A.

**Supplemental Figure S5.** The screen of optimal AbA concentration to inhibit the self-activation of *SlNAP2* and *SlEIL3*. AbA: Aureobasidin. A, Schematic representation of the SlNAP2 and SlEIL3 ligated to pGBKT7 vectors. B, The screen of optimal AbA concentration to inhibit the self-activation of *SlNAP2* and *SlEIL3*.

## Acknowledgments

The authors thank Jingquan Yu and Xiaojian Xia at Zhejiang University for their help in obtaining the mutants.

